# *Toxoplasma* Cathepsin Protease B and Aspartyl Protease 1 play recessive roles in endolysosomal protein digestion during infection

**DOI:** 10.1101/835405

**Authors:** Christian McDonald, David Smith, Manlio Di Cristina, Geetha Kannan, Zhicheng Dou, Vern B. Carruthers

## Abstract

The lysosome-like vacuolar compartment (VAC) is a major site of proteolysis in the intracellular parasite *Toxoplasma gondii*. Previous studies have shown that genetic ablation of a VAC-residing cysteine protease, cathepsin protease L (CPL), resulted in accumulation of undigested protein in the VAC and loss of parasite viability during the chronic stage of infection. However, since the maturation of another VAC localizing protease, cathepsin protease B (CPB), is dependent on CPL, it remained unknown whether these defects result directly from ablation of CPL or indirectly from a lack of CPB maturation. Likewise, although a previously described cathepsin D-like aspartyl protease 1 (ASP1) could also play a role in proteolysis, its definitive residence and function in the *Toxoplasma* endolysosomal system was not well defined. Here we demonstrate that CPB is not necessary for protein turnover in the VAC and that CPB deficient parasites have normal growth and viability in both the acute and chronic stages of infection. We also show that ASP1 depends on CPL for correct maturation and it resides in the *T. gondii* VAC where, similar to CPB, it plays a dispensable role in protein digestion. Taken together with previous work, our findings suggest that CPL is the dominant protease in a hierarchy of proteolytic enzymes within the VAC. This unusual lack of redundancy for CPL in *T. gondii* makes it a single exploitable target for disrupting chronic toxoplasmosis.

---

The apicomplexan parasite *Toxoplasma gondii* is an obligate intracellular protozoan that chronically infects ∼2 billion people worldwide (1). Toxoplasmosis can be fatal for those who are immunocompromised, such as AIDS or organ transplant patients (1, 2). In pregnant women, *Toxoplasma* can overcome barriers at the maternal-fetal interface and parasitize the developing fetus, which can cause a spontaneous abortion or neurological pathologies in the surviving offspring (3). Strikingly, most immunocompetent infected individuals are asymptomatic despite being persistently infected. Insufficient understanding of the underlying pathways that permit long-term infection precludes the development of viable treatment options to limit disease in at-risk individuals.

The endolysosomal pathway is a vital system in eukaryotic organisms. Proteases associated with these systems are of particular interest due to their role in various biological processes including catabolism and enzyme activation (2, 4, 5). The lysosomal cysteine proteases cathepsins L and B are ubiquitously expressed in eukaryotic cells and together are integral for proteolytic activity within lysosomes (6). Single knockout experiments in murine models only show a modest or indistinguishable phenotype for cathepsin L or B deficiency, respectively (7, 8). For example, mouse embryonic fibroblasts lacking cathepsin L present a non-lethal deficit in the turnover of autophagosomes (7). This indicates that the absence of one of these proteases can be overcome in mice. However, double-knockout experiments of both CPL and CPB had a profound effect on mouse health, including pronounced neurodegeneration, defects in motility and most significantly, lethality three weeks after birth (6, 9). In addition, mice deficient in the lysosomal aspartyl protease cathepsin D die within 4 weeks of birth (10). Cathepsin D also regulates activation of cathepsin B (11). Thus, whereas cathepsins L and B are individually dispensable in mammals, cathepsin D is critical for maintaining proteolytic activity and survival in mammals.

Lysosomal cathepsin proteases are expressed in multicopy families in some pathogenic protozoa such as *Plasmodium falciparum. P. falciparum* is one of the causative agents of malaria and, like many other protozoan parasites, relies on proteases for various aspects of its pathogenesis (2). *Plasmodium* obtains host haemoglobin and digests it in a specialized organelle known as the food vacuole (FV) to procure necessary nutrients (12, 13). Three cathepsin L-like cysteine proteases (falcipains) and four cathepsin D-like aspartyl proteases (plasmepsins) localize to the FV and are key players in haemoglobin degradation (14–16). As with the mammalian systems mentioned above, there exists functional redundancy between these proteases (13, 17, 18). Knockout of falcipain-2a in one study resulted in an accumulation of unprocessed haemoglobin in the FV; however, this was a transient phenotype and was subsequently not seen in later stages of the parasite life cycle presumably due to expression of falcipains 2b and 3 (17). Likewise, parasites lacking any one of the four plasmepsins were slower growing yet viable (18). Taken together, this suggests these proteases work cooperatively in the turnover of host haemoglobin.

Like its *Plasmodium* kin, *Toxoplasma* possesses a recently characterized lysosome-like organelle known as the vacuolar compartment (VAC), which houses proteases canonically associated with lysosomes (19–21). Acute stage parasites (tachyzoites) have been shown to ingest host cytosolic proteins for the duration of the parasite cell cycle, and the catabolism of these host-derived constituents occurs in the VAC (19, 22). In contrast to *Plasmodium, Toxoplasma* encodes single copies of cathepsin L (CPL), cathepsin B (CPB), and cathepsin D (aspartyl protease 1, ASP1) in its genome. Of the five cathepsin-like cysteine proteases encoded in the *Toxoplasma* genome, the VAC is the major site for CPB and CPL (23–25). Recent investigations of the VAC are uncovering how chronic stage parasites (bradyzoites) sustain themselves in the host indefinitely following encystation. First, genetic ablation of CPL has shown its contribution towards acute stage virulence and invasion. Second, cystogenic strains of *T. gondii* deficient in this protease accumulate large quantities of undigested material in the VAC of bradyzoites *in vitro*. Finally, the viability of CPL-deficient bradyzoites was starkly reduced *in vitro*, consistent with vastly reduced cyst burden *in vivo* (26). It remains unclear whether this is solely due to CPL disruption or due to CPB maturation being dependent on CPL (23). In other words, it remains unknown whether the phenotypes observed in CPL-deficient parasites are due to a lack of active CPB.

Of the seven aspartyl proteases identified in the *Toxoplasma* genome only one, ASP1, shares a clade with the *P. falciparium* plasmepsins (27, 28). This phylogenetic relationship may point to a digestive function within the *T. gondii* VAC, similar to plasmepsins in the FV. However, its definitive location and role in the endolysosomal system has not been established.

Although a critical requirement for protein turnover in the VAC in chronic stage *T. gondii* has previously been shown, key questions remain regarding the contribution of specific proteases to this degradation system. Therefore, in this study our aim was to define the role of *T. gondii* CPB and ASP1 in protein turnover in the VAC, delineate their position within a protease hierarchy and determine their contribution to infection.

## RESULTS

### Determining the extent of CPB’s role in tachyzoite growth and host-derived protein turnover

Maturation of *Toxoplasma* protease CPB has been shown to be dependent on another VAC-residing protease, CPL. Since CPL deficient parasites exhibit numerous phenotypes and do not produce mature CPB, it is unclear whether the phenotypes described are attributed to inactive CPB. To assess the role of CPB in *T. gondii*, we knocked out CPB in the cystogenic type II Prugniaud Δ*ku80* strain (29) (termed Pru hereafter) and confirmed this by PCR analysis (Supplemental Fig. S1), immunoblotting and immunofluorescence (Fig. 1A & B). Inclusion of the previously described PΔ*cpl* strain (26) in the immunoblot confirmed that maturation of CPB is dependent on expression of CPL, which is consistent with previous findings in a type I strain (23). Similarly, we found that CPB localizes to the VAC of tachyzoites (Fig. 1B), which is also consistent among strains (23). To test how tachyzoite replication was affected following CPB knockout, Pru, PΔ*cpb* and PΔ*cpl* tachyzoites were grown in human foreskin fibroblast (HFF) cells followed by the enumeration of parasites per parasitophorous vacuole (PV) to quantify replication. At 24 h post-infection, no measurable differences were seen between these strains; however, at 48 h post-infection, PΔ*cpl* tachyzoites had slower growth than Pru, whereas replication of PΔ*cpb* was indistinguishable from Pru (Fig. 1C). As mentioned above, ingestion of host-derived material has been shown in the VAC of acute stage parasites. Using a Chinese Hamster Ovarian (CHO)-K1 cell line inducibly expressing mCherry, we found that accumulation of host-derived mCherry was significantly higher in PΔ*cpl* tachyzoites compared to both Pru and PΔ*cpb* strains, whereas PΔ*cpb* parasites showed comparable accumulation to Pru (Fig. 1D). In all, these findings suggest that CPB has a limited role in tachyzoite VAC digestive function.

**Figure 1.**
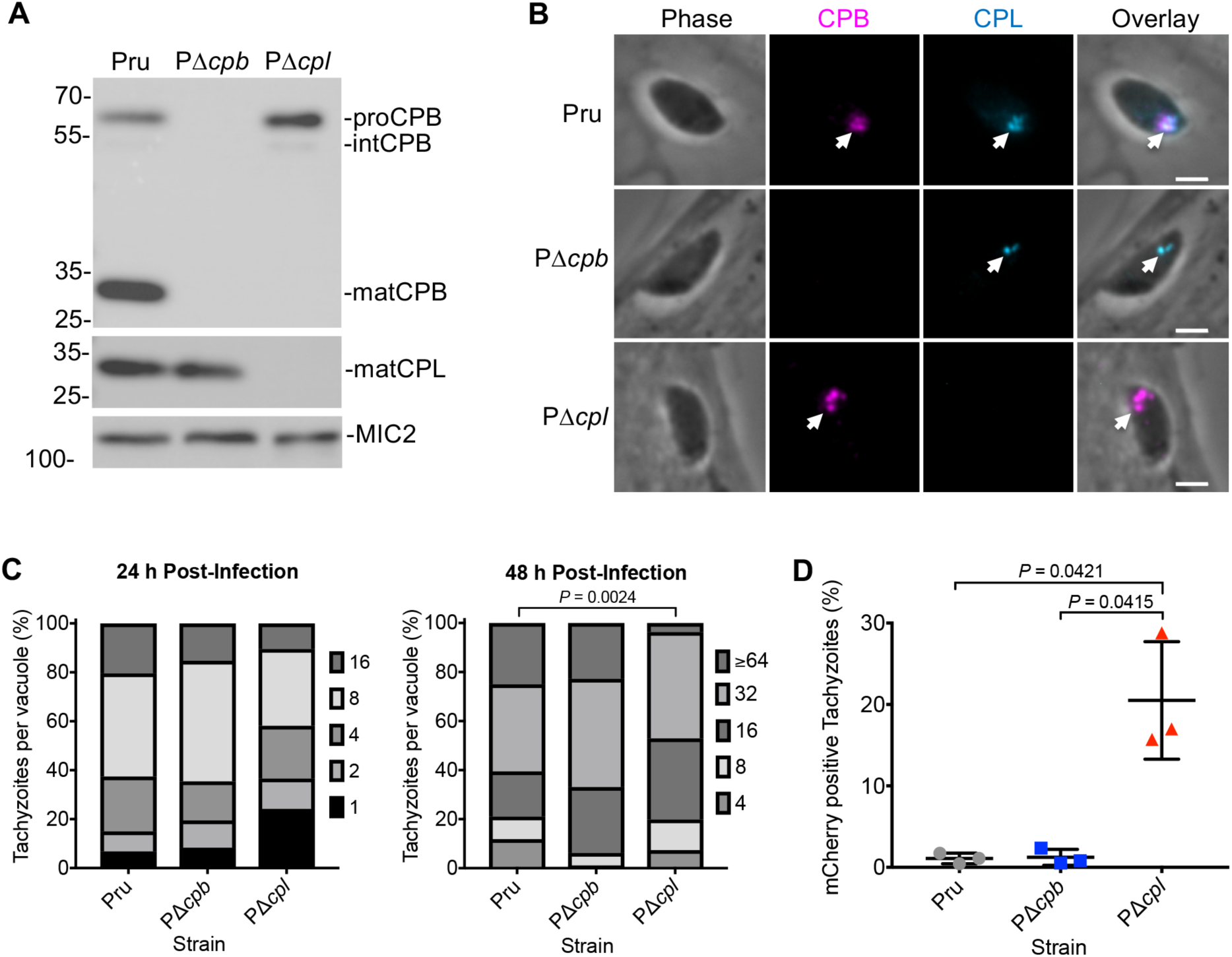
CPB does not play a major role in tachyzoite growth and protein turnover. *A*, Western blot of tachyzoite lysates probed for *T. gondii* CPL and CPB along with MIC2 as a loading control. Protein bands are labeled as proform (pro), intermediate (int) or mature (mat) form based on their molecular weights. *B*, Recently invaded intracellular tachyzoites were stained with anti-CPB (violet) and anti-CPL (blue). Arrowheads indicate labeling of the VAC. Scale bars, 2 µm. *C, T. gondii* tachyzoites were cultured in HFF monolayers and samples were collected at 24 and 48 h post-infection, fixed and stained with anti-SAG1 antibody, and the number of parasites per vacuole were quantified. Percentages of various replication stages for each strain are quantified. Results represent means from 2-3 biological replicates. The total number of parasitophorous vacuoles counted across 3 biological replicates was Pru: 147 and 152, PΔ*cpb:* 144 and 145, PΔ*cpl:* 153 and 136 for 24 and 48 h post-infection, respectively. Only statistically significant differences are represented in the figure. Statistical significance determined by unpaired two tailed Student’s *t* test performed on the mean number of parasites per vacuole for each strain across 3 biological replicates. *D*, Tachyzoite ingestion of mCherry assay quantification showing means ± standard deviation of 3 biological replicates. The data generated was analyzed using the following number of tachyzoites per replicate: Pru (*n*= 469, 464 and 231), PΔ*cpb* (*n*= 374, 613 and 213), PΔ*cpl* (*n*= 427, 299 and 216). All strains were compared and only significant differences are shown in the figure. Unpaired two-tailed *t* test with Welch’s corrections was performed on the mean of 3 biological replicates.

### CPB maturation depends on CPL activity in bradyzoites

We used a previously described series of strains (26) to explore the role of CPL activity in the maturation of itself and CPB in chronic stage bradyzoites. PΔ*cplS1CPL* and PΔ*cplB1CPL* express CPL exclusively in tachyzoites and bradyzoites, respectively. PΔ*cplB1CPL*^*^ exclusively expresses a catalytically inactive form of CPL in bradyzoites. Using immunoblotting, we found CPL catalytic activity is necessary for normal self-maturation based on observing a pseudomature (pmatCPL) species in PΔ*cplB1CPL*^*^ bradyzoites (Fig. 2A). These results are consistent with previous findings in tachyzoites (23). Also, when probed for CPB, immunoblots displayed bands for mature CPB only in strains that express active CPL, namely Pru and PΔ*cplB1CPL* bradyzoites (Fig. 2B). Immature CPB was observed in Δ*cplB1CPL*^*^, thus confirming that CPL activity is necessary for maturation of CPB in bradyzoites.

**Figure 2.**
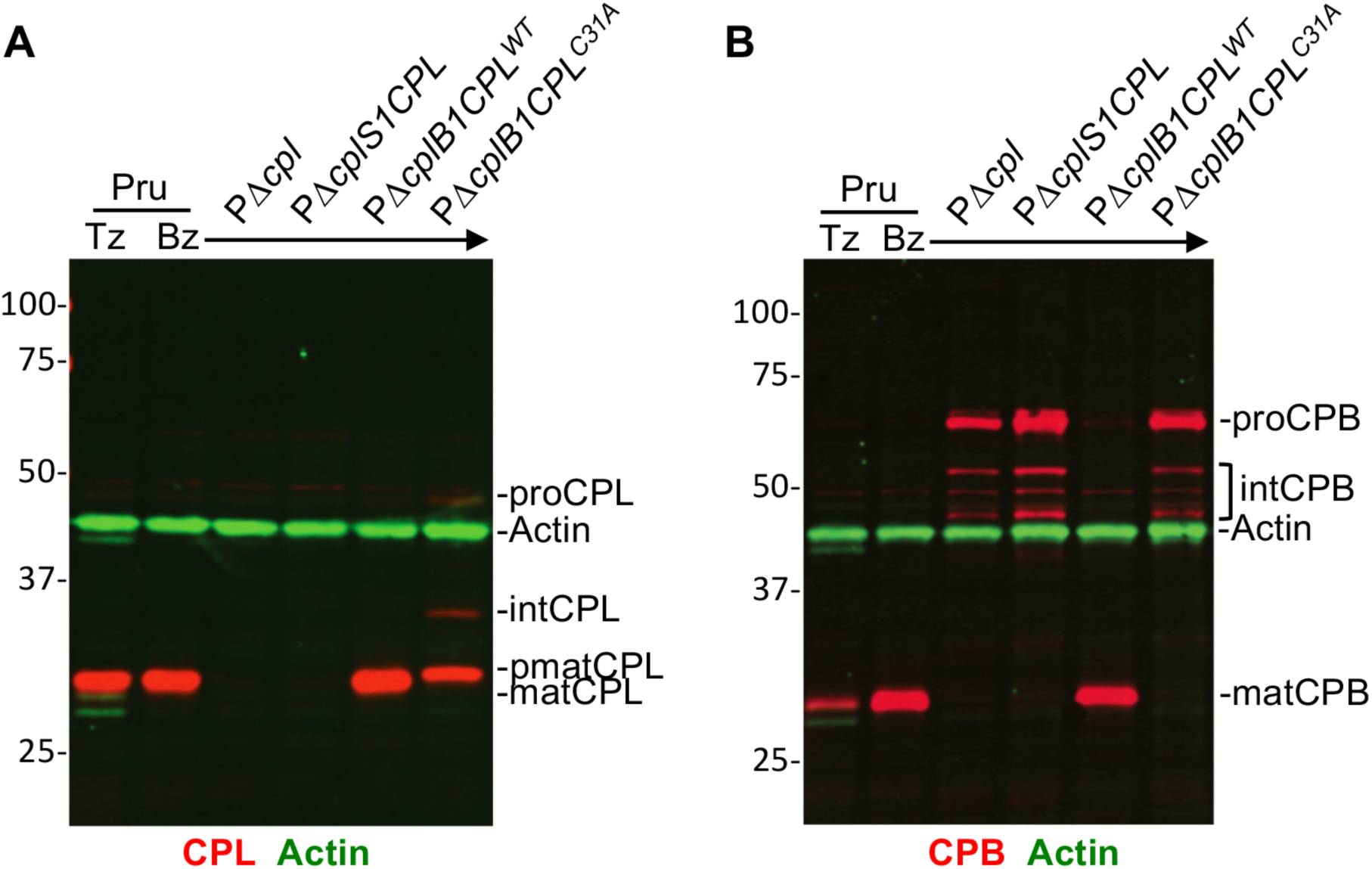
*T. gondii* CPB maturation is dependent on the expression of active CPL. *A & B*, Li-COR western blots of tachyzoite and bradyzoite cell lysates probed for *T. gondii* CPL (red-panel A), CPB (red-panel B) and actin (green-panels A & B) as the loading control. Protein bands are labeled as proform (pro), intermediate (int), pseudomature (pmat) and mature (mat) form based on their molecular weights.

### CPB is not necessary for turnover of autophagic material in bradyzoites

We have previously demonstrated that CPL-deficiency in *T. gondii* leads to the accumulation of autophagic material in the VAC in bradyzoites (26). However, due to CPB maturation being dependent on the proteolytic activity of CPL, parasites deficient in CPL will also be deficient in active CPB. Therefore, it remained a possibility that this accumulation was actually due to a lack of active CPB in the VAC. To address this we first used surface antigen 1 (SAG1) as a tachyzoite stage-specific marker as well as GFP, which is expressed following bradyzoite differentiation (29) and found that differentiation was similar among the Pru, PΔ*cpb*, and PΔ*cpl* strains (Fig. 3A). Next, consistent with our observations of CPB and CPL colocalization in tachyzoites, these two proteases were also found to colocalize in bradyzoite cysts (Fig. 3B). To determine the extent to which CPB independently contributed to the turnover of autophagic material, we assessed the accumulation of dense puncta indicative of undigested autophagosomes in PΔ*cpb* bradyzoites and compared this to Pru and PΔ*cpl* bradyzoites. Our results showed that after 7 and 14 days of differentiation, punctate structures were larger in PΔ*cpl* bradyzoites compared Pru (Fig. 3C & D). However, no difference in puncta size was seen in PΔ*cpb* bradyzoites compared to Pru. Collectively, our findings suggest that CPB does not play a major digestive role in the endolysosomal system of *Toxoplasma* during acute or chronic infection *in vitro*.

**Figure 3.**
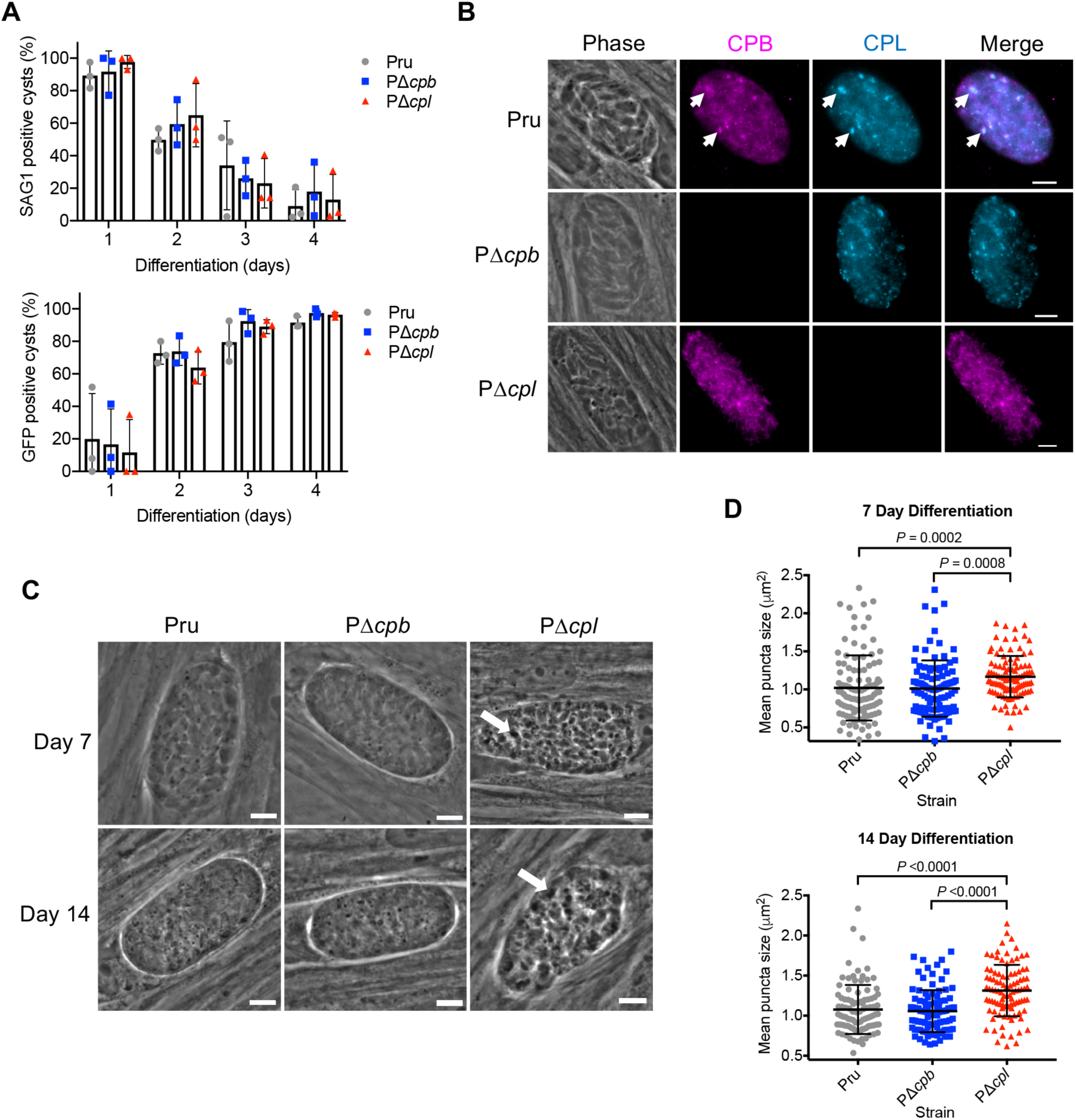
CPL, not CPB, is the major protease necessary for the turnover of autophagosomes in the VAC in bradyzoites. *A*, The rate of parasite differentiation from the tachyzoite stage to the bradyzoite stage was determined *in vitro* by assessment of the tachyzoite-specific antigen SAG1 and GFP under the control of a bradyzoite LDH2 promoter. Infected monolayers were cultured for the indicated days, fixed, probed for SAG1 and quantified. Bars indicate means ± standard deviation of 3 biological replicates. Experiments analyzed for Pru 105, 106, 153 and 151 cysts, for PΔ*cpb*117, 123, 149 and 110 cysts and for, PΔ*cpl* 75, 108, 155 and 130 cysts on days 1,2,3 and 4, respectively. All strains were compared and only statistical significance is shown in the figure. Unpaired two-tailed *t* test with Welch’s corrections was performed on the means of 3 biological replicates. *B*, Immunofluorescence localization of CPB and CPL in bradyzoites. Parasite strains were differentiated for 7 days, fixed and stained with anti-CPB (red) and anti-CPL (blue). Scale bars, 5 µm. *C*, Phase-contrast microscopy was used to image bradyzoites *in vitro* following 7- and 14-day cultures. Enlarged dark puncta seen in PΔ*cpl* bradyzoites are indicative of defective protein degradation in the VAC (arrows). Scale bars, 5 µm. *D*, The size of the puncta in bradyzoites was measured. The following numbers of cysts across 3 biological replicates were used to analyse each strain: Pru (Day 7: 109 cysts; Day 14: 98 cysts), PΔ*cpb* (Day 7: 106 cysts; Day 14: 102 cysts), PΔ*cpl* (Day 7: 105 cysts; Day 14: 97 cysts). Lines represent means ± standard deviation. For this data set, ROUT with a Q value of 0.1% was used to identify and remove 1 outlier each for Pru and PΔ*cpl* for 14-days post-differentiation. A Kruskal-Wallis test with Dunn’s multiple comparisons was performed on the means across 3 biological replicates. All strains were compared and only significant differences are shown.

### ASP1 resides in the Toxoplasma VAC and is dependent on CPL for normal maturation

Unlike the related apicomplexan, *Plasmodium, Toxoplasma* possesses one group A aspartyl protease, ASP1 (27). *T. gondii* ASP1 was shown to have punctate localization that extensively changes during tachyzoite replication in daughter cells and was predicted to be compartmentalized (27). The *Toxoplasma* VAC displays similar fragmentation during replication (21) and we therefore hypothesized that ASP1 resides within the VAC. To address this definitively, we generated antibodies to full length recombinant ASP1 and created an ASP1 deficient mutant in the RH strain background (RΔ*asp1*) along with genetically complementing the mutant (RΔ*asp1ASP1*)(Supplemental Fig. S2). Immunoblots displayed mature ASP1 in parental strain RH and RΔ*asp1ASP1* complement strain, but not in RΔ*asp1* (Fig. 4A). Note that despite affinity purifying the antibody on recombinant ASP1, several non-specific bands were observed in the RΔ*asp1* lysate including one that nearly co-migrates with mature ASP1. PCR analysis (Supplemental Fig. S2) and immunofluorescence staining confirmed the absence of ASP1 expression in RΔ*asp1* tachyzoites. Immunofluorescence staining also revealed colocalization of ASP1 with CPL in RH and RΔ*asp1ASP1* (Fig. 4B).

**Figure 4.**
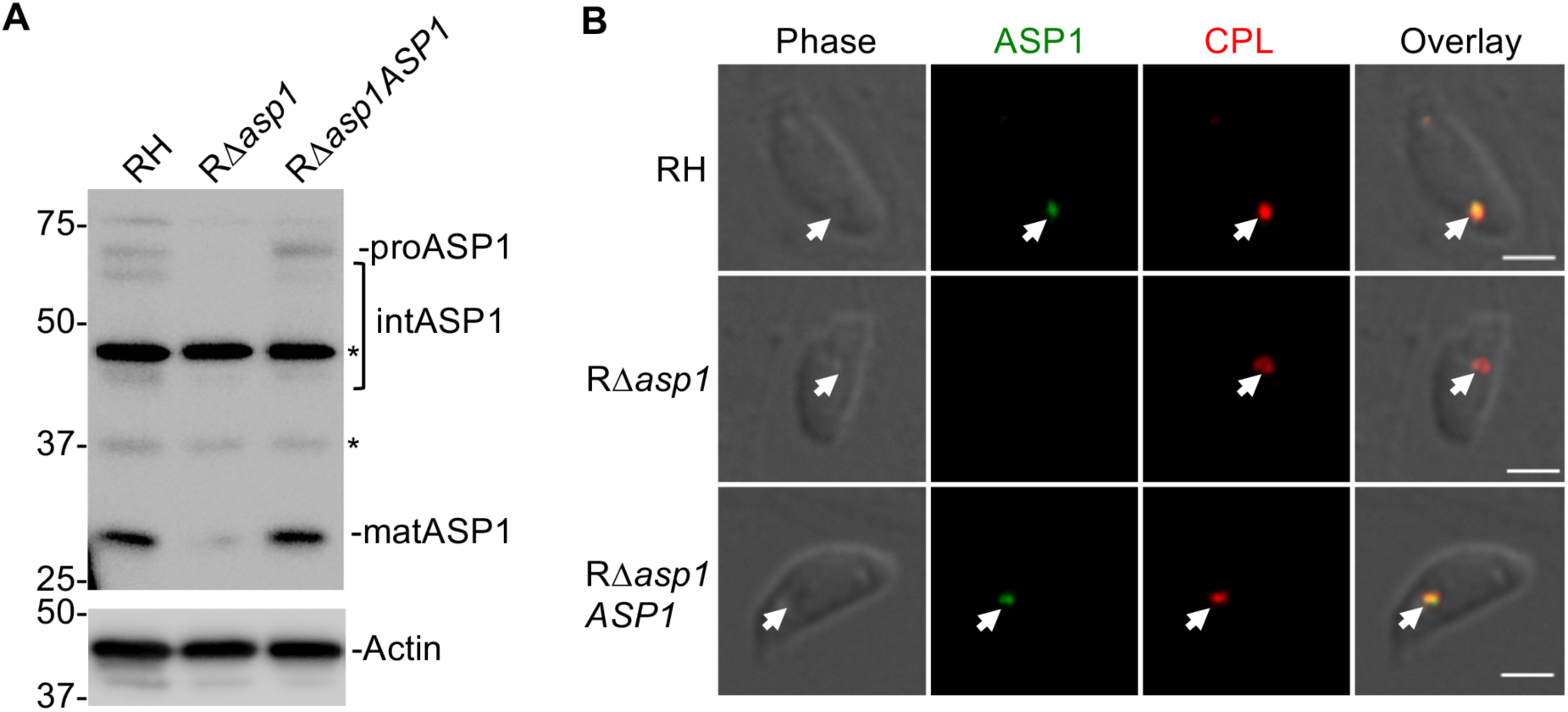
*T. gondii* ASP1, a cathepsin D ortholog that localizes to the VAC. *A*, Immunoblot detection of *T. gondii* ASP1 in tachyzoites. Protein bands are labeled as proform (pro), intermediate (int) and mature (mat) form based on their molecular weights. Polypeptides marked with asterisks are non-specific bands. *B*, Immunofluorescence assay of tachyzoites after fixation, staining and probing for *T. gondii* ASP1 (green) and CPL (red). Arrowheads indicate co-localization of ASP1 with CPL in the VAC. We did not observe ASP1 staining in RΔ*asp1* parasites. Scale bars = 2 µm.

Since CPL expression is required for maturation of CPB, we reasoned that CPL might also contribute to the maturation of ASP1. Accordingly, immunoblotting of RΔ*cpl* with anti-ASP1 revealed accumulation of proASP1 and intermediate species of ASP1 (Fig. 5A), suggesting a role for CPL expression in ASP1 maturation. Note that the non-specific band nearly comigrating with mature ASP1 confounded interpretation of how much mature ASP1 is present in RΔ*cpl* tachyzoites. We also observed a corresponding increase in the staining for ASP1 and CPB in RΔ*cpl* tachyzoites (Fig. 5B), indicating accumulation of immature species of these proteases in the absence of CPL expression. Collectively, our findings suggest that ASP1 resides in the VAC and that its maturation is at least partially dependent upon expression of CPL.

**Figure 5.**
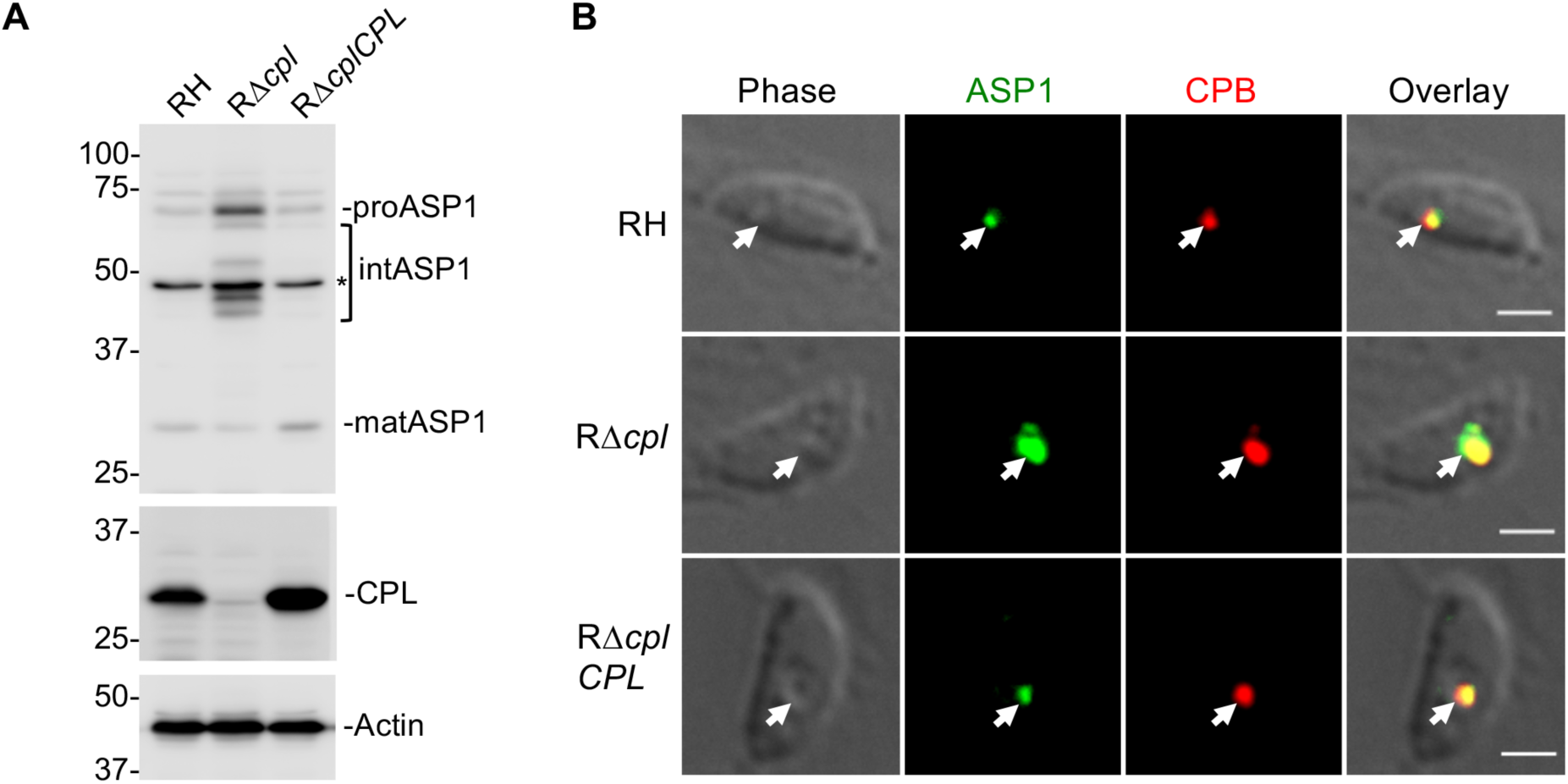
*T. gondii* immature ASP1 accumulates in tachyzoites lacking active CPL in the VAC. *A*, Immunoblot of tachyzoite lysates probed for ASP1 and CPL along with Actin as a loading control. For ASP1, based on molecular weights, protein bands are labeled as proform (pro), intermediate (int) and mature (mat) form. *B*, Immunofluorescence assay to detect *T. gondii* ASP1 (green) and CPB (red) in RH, RΔ*cpl* and RΔ*cplCPL* tachyzoites. Arrowheads show localization of ASP1 with CPB in the VAC. Scale bars = 2 µm.

### ASP1 is not required for tachyzoite replication or digestion of host-derived protein in the VAC

To further investigate the maturation and function of ASP1 in infection, we generated Pru parasites lacking ASP1 alone (PΔ*asp1*) or lacking ASP1 and CPL (PΔ*asp1*Δ*cpl*) (Supplemental Fig. S3). Immunoblotting of tachyzoite (Fig. 6A, left panel) or bradyzoite (Fig. 6A, right panel) corroborated a role for CPL in ASP1 maturation. Also, because the non-specific band that co-migrated with mature ASP1 is absent in bradyzoites, it is apparent that maturation of ASP1 is strongly dependent on expression of CPL in bradyzoites. The increased expression of both ASP1 and CPL in bradyzoites compared to tachyzoites is also evident (Fig. 6A, right). To assess the role of ASP1 in tachyzoite replication, parasites were allowed to infect HFF monolayers for 24 and 48 h. Enumeration of the parasites per PV displayed significant differences only after 48 h post infection in PΔ*cpl* and PΔ*asp1*Δ*cpl* strains (Fig. 6B). Interestingly, PΔ*asp1*Δ*cpl* parasites did not replicate slower than Δ*cpl* parasites, implying that the absence of CPL is principally responsible for the replication phenotype. Next we tested for a possible role for ASP1 in digestion of host-derived protein in the VAC. Following the incubation of parasites with mCherry expressing CHO-K1 cells, PΔ*asp1* failed to accumulate host-derived mCherry whereas accumulation was seen in PΔ*cpl* and PΔ*cpl*Δ*asp1* tachyzoites (Fig. 6C). These results mirror what was seen in CPB and indicate that neither CPB nor ASP1 are major contributors to tachyzoite growth or endolysosomal digestive function, whereas CPL has a dominant role.

**Figure 6.**
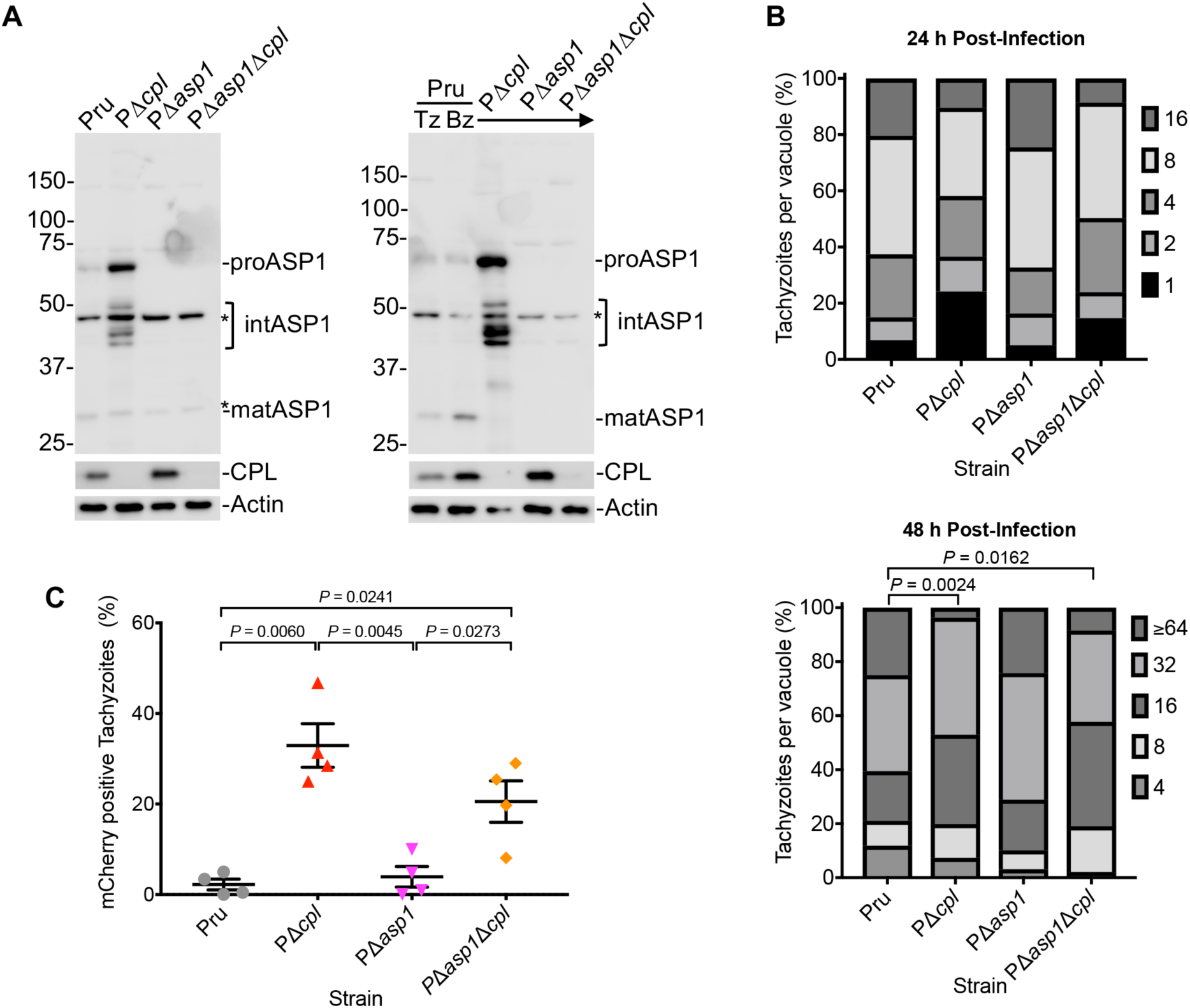
ASP1 is dispensable for tachyzoite replication or ingestion. *A*, Parasite lysates of tachyzoites and bradyzoites were immunoblotted with antibodies against *T. gondii* ASP1 and CPL along with Actin as a loading control. For ASP1, based on molecular weights, protein bands are labeled as proform (pro), intermediate (int) and mature (mat) form. *B, T. gondii* tachyzoites were cultured in HFF monolayers and samples were collected at 24 and 48 h post-infection, fixed and stained with anti-SAG1 antibody and the number of parasites per vacuole were quantified. Percentages of various replication stages for each strain are quantified. The total number of parasitophorous vacuoles counted across 2 biological replicates was Pru: 147 and 152, PΔ*cpl:* 153 and 136, PΔ*asp1*: 159 and 128, PΔ*asp1*Δ*cpl*: 163 and 142, for 24 and 48 h post-infection, respectively. Only statistically significant differences are represented in the figure. Statistical significance determined by unpaired two tailed Student’s *t* test performed on the mean number of parasites per vacuole for each strain across 3 biological replicates. *C*, Tachyzoite ingestion of mCherry assay quantitation showing means ± standard deviation of 4 biological replicates. The data generated was analyzed using the following number of tachyzoites per replicate: Pru (*n*= 205, 219, 223 and 207), PΔ*cpl* (*n*= 224, 242, 224 and 223), PΔ*asp1* (*n*= 218, 220, 220 and 210) and PΔ*asp1*Δ*cpl* (*n*= 203, 235, 228 and 246). All strains were compared and only significant differences are shown in the figure. Unpaired two-tailed *t* test with Welch’s corrections was performed to determine statistical significance.

### ASP1 plays a dispensable role in proteolytic turnover of autophagic material

It was also important to establish whether ASP1 could have a functional overlap in the turnover of autophagic material in bradyzoites. Therefore, similar to that for PΔ*cpb* parasites, we compared differentiation and size of puncta in PΔ*asp1* and PΔ*asp1*Δ*cpl* bradyzoites to those of Pru and PΔ*cpl* bradyzoites after 7 and 14 days of differentiation. Differentiation was indistinguishable among the strains based on the decrease in SAG1 positive vacuoles and consequential increase of GFP positive cysts (Fig. 7A). Also, automated measurement of puncta size indicated normal turnover of autophagosomes in PΔ*asp1* bradyzoites, which is distinct from the enlarged puncta observed in PΔ*cpl* and PΔ*asp1*Δ*cpl* bradyzoites (Fig. 7B & C). Taken together, our results suggest that CPB and ASP1 do not play a major role in the proteolytic turnover of autophagic material in the VAC in bradyzoites, whereas CPL has a dominant role in this activity.

**Figure 7.**
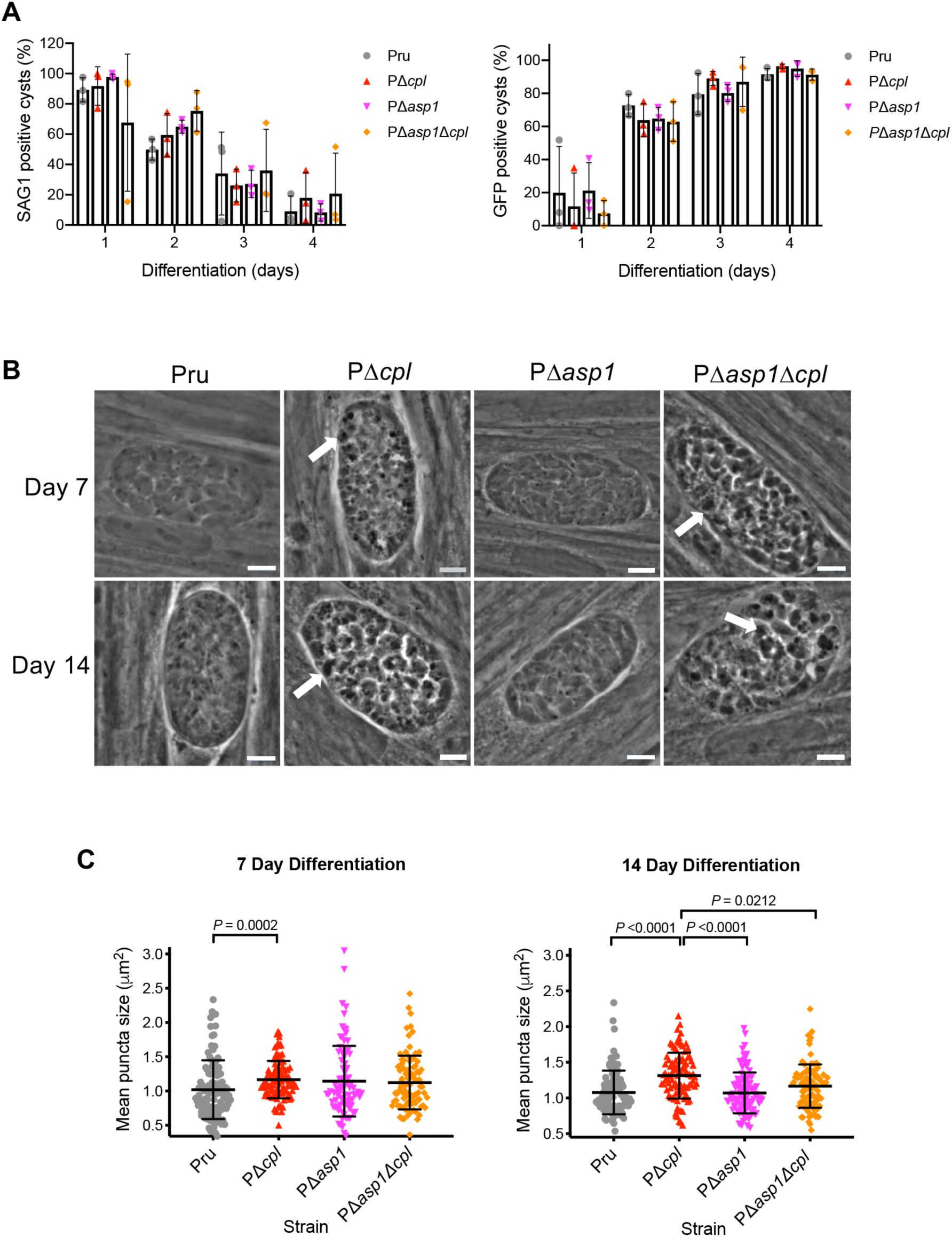
*T. gondii* ASP1 has a limited role in protein turnover of autophagosomes. *A*, The rate of parasite differentiation from the tachyzoite stage to the bradyzoite stage was determined *in vitro* by assessment of the tachyzoite-specific antigen SAG1 and GFP under the control of a bradyzoite LDH2 promoter. Infected monolayers were cultured for the indicated days, fixed, probed for SAG1 and quantified. Bars indicate means ± standard deviation of 3 biological replicates. Experiments analyzed for Pru 105, 106, 153 and 151 cysts, for PΔ*cpl* 75, 108, 155 and 130 cysts, for PΔ*asp1* 119, 124, 189 and 146 cysts and for PΔ*asp1*Δ*cpl* 78, 110, 112 and 161 cysts on days 1, 2, 3 and 4, respectively. All strains were compared and only statistical significance is shown on the figure. Unpaired two-tailed *t* test with Welch’s corrections was performed on the mean across 3 biological replicates. *B*, Phase-contrast microscopy was used to image bradyzoites *in vitro* following 7- and 14-day culture under differentiation conditions. Enlarged dark puncta indicative of defective protein degradation in VACs are visible in PΔ*cpl* bradyzoites and PΔ*asp1*Δ*cpl* cysts (arrows). Scale bars, 5µm. *C*, The size of the puncta in bradyzoites was measured. The following numbers of cysts across 3 biological replicates were used to analyse each strain: Pru (Day 7: 109 cysts; Day 14: 98 cysts), PΔ*cpl* (Day 7: 105 cysts; Day 14: 97 cysts), PΔ*asp1* (Day 7: 85 cysts; Day 14: 97 cysts), PΔ*asp1*Δ*cpl* (Day 7: 90 cysts; Day 14: 97 cysts). Lines represent means ± standard deviation. For this data set, ROUT with a Q value of 0.1% was used to identify and remove 1 outlier from PΔ*asp1* for 7-days post-differentiation and 1 outlier each for Pru and PΔ*cpl* for 14-days post-differentiation. A Kruskal-Wallis test with Dunn’s multiple comparisons was performed on the mean across 3 biological replicates. All strains were compared and only significant differences are shown.

### CPL plays a principal role in bradyzoite viability

To quantify the extent to which CPL, CPB and ASP1 contribute to bradyzoite viability, we measured the ability of bradyzoites lacking these proteases to invade, differentiate to tachyzoites and replicate within HFF monolayers, which if left undisturbed results in the formation of plaques (Fig. 8A). Because proteolytic deficiency in the VAC is associated with accumulation of puncta and no increases in puncta were observed for PΔ*cpb* or PΔ*asp1* between 7 and 14 days of differentiation, we only measured bradyzoite viability at 7 days of differentiation. As expected, PΔ*cpl* bradyzoites showed a marked loss of viability based on plaque formation (Fig. 8B & C). By contrast, viability of PΔ*cpb* and PΔ*asp1* bradyzoites were no different from that of Pru. Moreover, the decreased viability of PΔ*asp1*Δ*cpl* bradyzoites was no greater than that of PΔ*cpl* bradyzoites as compared to Pru bradyzoites. Together, these findings suggest that whereas CPL is a key enzyme in bradyzoites, CPB and ASP1 do not play a substantial role in bradyzoite survival.

**Figure 8.**
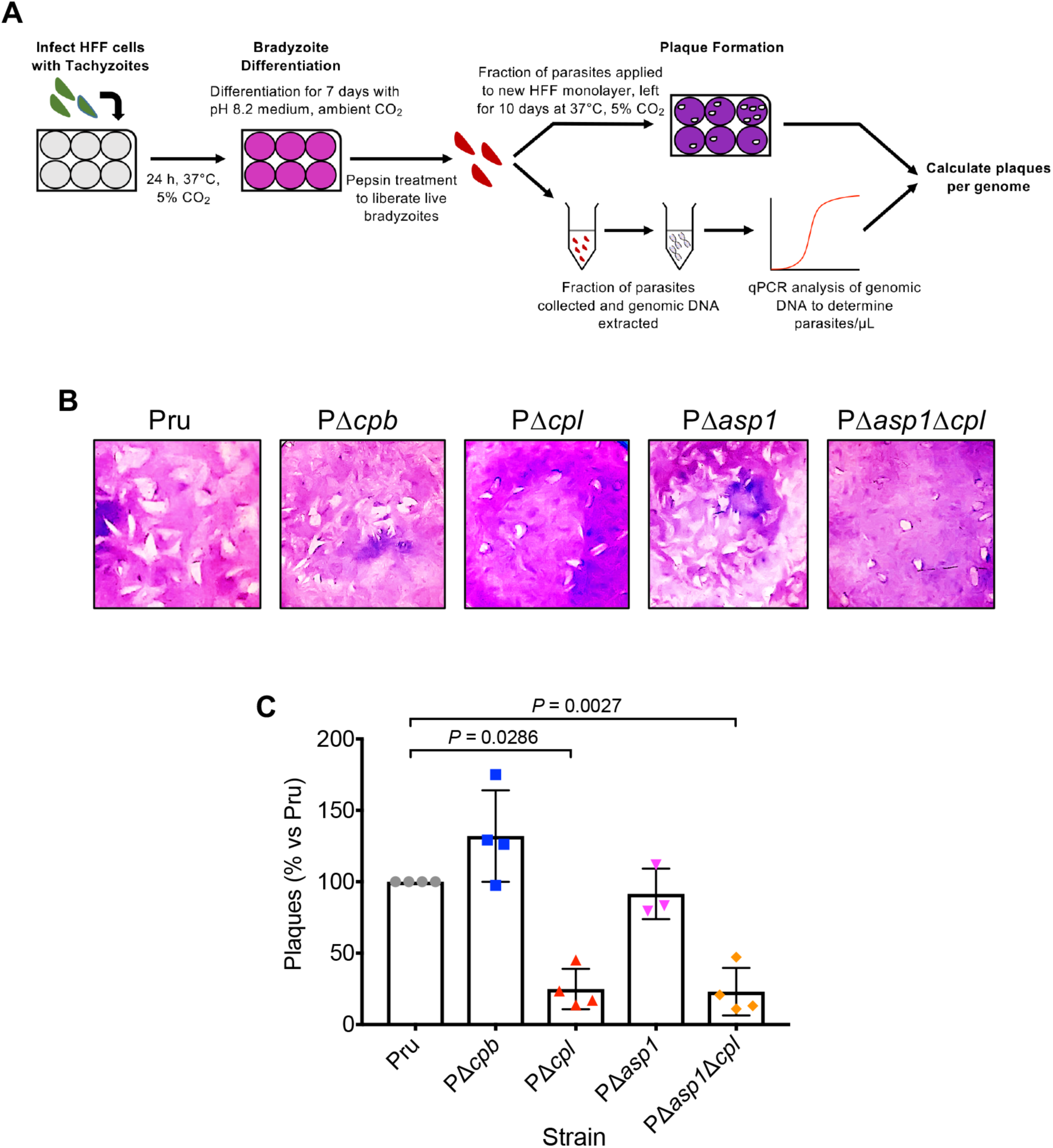
CPL is a central protease in bradyzoite viability. *A*, Schematic illustration of the workflow used to examine bradyzoite viability. Experiments conducted were used to generate the data collected in panels B & C. Details are provided in the Experimental Procedures Section. *B*, Plaque assays were performed on bradyzoites to determine the viability of each strain, based on their ability to undergo the lytic cycle following forced encystation. Plaque zones were visualized following crystal-violet staining of the HFF monolayer. *C*, Quantitation of plaque assay described in panel B. Quantitative PCR analysis was used to determine the number of parasites applied to fresh HFF monolayers for the bradyzoite plaque assay, allowing for the number of plaques per genome to be calculated. Data was normalized to the parental strain (Pru) and shown as a percentage. Bars indicate means ± standard deviation of 3 to 4 biological replicates. Only significant differences between Pru and subsequent strains is indicated in the figure. Mann-Whitney test was used to assess statistical significance.

## DISCUSSION

*Toxoplasma gondii* cathepsin L (CPL) was previously shown to have a role in protein turnover within the parasite VAC (19, 26). CPL is also necessary for the maturation of *T. gondii* CPB (23). Extending these findings herein, we demonstrate that CPB and CPL co-localize in chronic stage bradyzoites. Accordingly, it was necessary to assess whether a block in protein turnover in the VAC of CPL-deficient parasites was actually due to a lack of active CPB. Using ingestion assays to assess their contribution to protein digestion in the VAC in tachyzoites, we found that those lacking the *cpb* gene showed accumulation of host-derived mCherry comparable to the Pru parental strain. Similarly, we found comparable puncta in Δ*cpb* and WT bradyzoites, suggesting that the observed effect of protein accumulation in the VAC of CPL-deficient tachyzoites and bradyzoites is not due to a defect in CPB maturation.

Interestingly, we also found that an aspartyl protease (ASP1), which also occupies the VAC, was not necessary for protein degradation in the VAC or parasite viability. Further, although ASP1 abundance increases in CPL-deficient parasites, this is apparently due to a lack of its maturation rather than a compensatory mechanism. That parasites lacking both ASP1 and CPL are no worse off than those lacking CPL alone essentially rules out compensation by ASP1 in the absence of CPL. Transcriptomic datasets show that *CPL, CPB* and *ASP1* are each up-regulated in chronic stage bradyzoites (30) compared to acute stage tachyzoites, which is further supported by our immunoblotting results for CPB and ASP1 in tachyzoites and bradyzoites. Although we found no evidence to suggest CPB and ASP1 are necessary for the turnover of proteins in the VAC in bradyzoites, their increased expression in bradyzoites suggests they might have another role in chronic infection. Despite this, we found that both CPB and ASP1 are not essential for bradyzoite survival. This is in stark contrast to CPL, reinforcing the notion that CPL is the major protease of the VAC in *T. gondii* bradyzoites.

The non-redundant functions between CPL with other VAC proteases in *T. gondii* are unlike what has been seen in other eukaryotic organisms previously described. In some other systems cathepsin L is dispensable. For example, in murine models it is only when cathepsin B and L are both deleted from the genome that a strong detrimental phenotype is observed (6, 9).

Cathepsin D is the major aspartyl protease in the mammalian endolysosomal system and is essential for mouse survival (10). This is in contrast to what has been observed in apicomplexans (previously in *P. falciparum* and reported here in *T. gondii*), whereby no single aspartyl protease resident within lysosome-like compartments has been shown to be essential to parasite fitness (18). Although not essential for survival, a quadruple knockout of all four plasmepsins in *P. falciparum* resulted in a slower growth rate, delayed schizont maturation and reduced hemozoin formation (31, 32). This differs to our findings in the related parasite *T. gondii* whereby no defect was observed in stage differentiation and protein turnover in Δ*asp1* parasites. It has also been shown here and elsewhere that ASP1 is dispensable for tachyzoite growth (28).

Limiting parasite access to essential nutrients by blocking protein turnover has become a major focus of anti-parasite drug development (33–35). In *P. falciparum*, the hydrolysis of hemoglobin involves the cooperative action of two classes of proteases. In addition to their direct role in protein degradation, falcipains have an indirect role via activation of plasmepsins; however, autoprocessing of plasmepsins has also been observed following falcipain disruption (15). This has implications for the development of inhibitors against *P. falciparum* proteases involved in protein degradation in the FV, as they will have to block the function of multiple cathepsins simultaneously. Our work reveals a salient aspect of the *Toxoplasma* endolysosomal system in that, unlike *Plasmodium*, there exists an opportunity to target a single non-redundant protease, CPL.

*T. gondii* chronically infects one third of the worldwide population (1). Despite drug development against the infection being an active area of research, there is currently no effective treatment for chronic toxoplasmosis. Although previous work has focused on developing inhibitors against more than one cathepsin protease in *T. gondii* (36), it is now clear that CPL can be the focus of future efforts targeting disruption of VAC proteolysis. Of particular favor is that the development of compounds that specifically inhibit *T. gondii* CPL might not have severe side-effects resulting from inhibition of host cathepsin L (34).

## EXPERIMENTAL PROCEDURES

### Parasite cell culture and differentiation into bradyzoites

Throughout this investigation, Human Foreskin Fibroblast (HFF) cells were used to propagate parasite strains in Dulbecco’s modified Eagle Medium (DMEM) supplemented with 10% (v/v) cosmic calf serum (CCS) or fetal bovine serum (FBS).

For all bradyzoite conversion, tachyzoites were mechanically lysed by scraping infected HFF monolayers that were then passed through a 20 and a 23-gauge syringe and a 3 μm filter. Filtered parasites were then counted and allowed to infect fresh monolayers of HFF cells for 24 h. Bradyzoite differentiation was induced using alkaline pH medium and ambient CO_2_ (37–39). Briefly, 24 h after parasites were applied to HFF monolayers, DMEM media was replaced for an alkaline differentiation media (RPMI without NaHCO_3_, 50 mM HEPES, pen/strep, 3% FBS, pH 8.25). Differentiation media was replaced daily.

### Generation of transgenic T. gondii strains

Strategy, vectors and repair templates employed to create all the transgenic strains used in this study are described in the supplementary materials. Transfection of parasites was performed as described in (40). Briefly, 2 × 10^7^ tachyzoites were harvested, washed twice in cytomix [120 mM KCl, 0.15 mM CaCl_2_, 10 mM K_2_HPO_4_-KH_2_PO_4_ (pH 7.6), 25 mM HEPES (pH 7.6), 2 mM EGTA, 5 mM MgCl_2_, 2 mM ATP, 5 mM glutathione] and resuspended in 700 µl of cytomix containing either 5 µg of PCR repair template or 50 µg of plasmid. The entire mixture was then transferred to an electroporation cuvette (4-mm gap) and exposed to an electric pulse with an electroporator (BTX 600) set at 2.0 kV and 48. Electroporated parasites were transferred to fresh HFF monolayers and subjected to drug selection. Clones were isolated by limiting dilution.

### Expression and purification of recombinant pro-form ASP1

A ∼1.5-kb DNA fragment encoding the recombinant pro-form of ASP1 (rproASP1) was PCR-amplified from a *T. gondii* RH cDNA library using Q5^®^ High-Fidelity DNA polymerase (NEB) and cloned into a pET22b vector with a 6xHis epitope tag fused to the N-terminal end of ASP1. The resulting pET22b-6xHis-ASP1 expression construct was transformed into the *E. coli* protein expression strain, ER2566 (NEB), induced with 1 mM isopropyl 1-thio-β-D-galactopyranoside (IPTG) at 37°C for 4 h and purified using a standard purification procedure as described previously (25). The purified denatured ASP1 protein was refolded in PBS overnight by dialysis. Insoluble protein was removed by centrifugation at 20,000*g* for 10 min and the soluble fraction was concentrated using an Amicon^®^ Ultra 30-kDa cutoff concentrator (Millipore) and stored at - 20°C for future use.

### Immunization and affinity purification of anti-ASP1

Two New Zealand white rabbits were injected with 200 µg of purified recombinant proASP1 in Freund’s complete adjuvant and boosted 3 times at days 14, 21, and 56 with 50 µg of recombinant protein in Freund’s incomplete adjuvant (Cocalico Biologicals, Inc). The final sera were collected and further affinity purified. Purified recombinant proASP1 was crosslinked to 0.5 ml of AminoLink Coupling Resin (Thermo Fisher) and affinity purified anti-ASP1 antibodies were eluted with 0.2 M Glycine, pH 3.0. The elution fractions containing the antibodies were pooled, buffer-exchanged, and concentrated by using Amicon Ultra 30-kDconcentrators.

### Immunoblotting

*Toxoplasma gondii* parasites were purified as described above, counted using a hemocytometer and pelleted by centrifugation at 1800*g* for 10 min. Parasite pellets were resuspended in SDS-PAGE sample buffer containing 5 mM DTT and heated at 95°C for 5 min. Parasite proteins were separated by SDS-PAGE on 15% polyacrylamide gels and then transferred to an Immobilin-P PVDF membrane using semidry electroblotting. Blots were blocked using 0.1% (v/v) Tween 20 in PBS (PBST) containing 5% (w/v) nonfat dried milk. Blots were probed with the primary antibodies rabbit anti-CPL (1:8,000), mouse anti-CPB (1:2,000), rabbit anti-CPL (1:8,000), affinity purified rabbit anti-ASP1 (1:2,500) and mouse anti-MIC2 (1:5,000). Secondary horse radish peroxidase conjugated goat-anti mouse/rabbit IgG (Jackson Immunoresearch) diluted 1:10,000 in PBST, was applied. Bound antibody was visualised with the chemiluminescence substrate SuperSignal West Pico (Thermo Scientific) and imaged using a G:BOX Chemi XRQ imager (Syngene). Fluorescence immunoblots were as above except they were blocked in 50 mM Tris-HCl, pH 7.4/1.25% (v/v) fish gelatin/150 mM NaCl for 30 min and secondaries antibodies were IRDye 800CW conjugated goat anti-rabbit IgG conjugated and IRDye 680RD conjugated goat-anti mouse IgG conjugated. Blots were air-dried before imaged by LI-COR Odyssey CLx instrument.

### Immunofluorescence Assay

To confirm protease localization in tachyzoites, indirect immunofluorescence was performed. Parasites were allowed to infect HFF cells for 30 min in 8-well chamber slides at 37°C, 5% CO_2_. Infected cells were then fixed using 4% (v/v) methanol-free formaldehyde in PBS. Cells were permeabilized with 0.1% (v/v) Triton X-100 for 10 min., followed by 20 min of blocking with PBS containing 10% FBS. Blocked monolayers were incubated at RT for 40 min with the primary antibodies mouse anti-CPB (1:200), rabbit anti-CPL (1:500) or affinity purified rabbit anti-ASP1 (1:200). Following washes in PBS/1%FBS/1% normal goat serum, slides were probed with secondary fluorophore-conjugated antibodies (Jackson Immunoresearch), mounted using Mowiol, and imaged on a Zeiss Axio inverted fluorescence microscope equipped with a Zeiss axiocam 305 scMOS digital camera.

For indirect immunofluorescence on bradyzoites, freshly lysed parasites were placed onto an HFF cell monolayer and left at 37°C, 5% CO_2_ overnight. Invaded parasites were subsequently stimulated to differentiate into bradyzoites by culturing in alkaline differentiation media and incubating at 37°C, 0% CO_2_ for 7 days. Bradyzoite cultures were fixed, probed with antibodies and mounted as described above.

### Parasite ingestion assay

To assess the degradation of proteinaceous material within the lysosome-like VAC in tachyzoites, the previously described ingestion assay was used (41). Briefly, modified CHO-K1 cells were induced to express cytosolic mCherry following treatment with doxycycline (2 µg/mL) for 5 days prior to infection. On the sixth day, freshly harvested parasites were allowed to infect and ingest cytosolic mCherry from doxycycline-treated CHO-K1 cells for 4 h. After incubating, parasites were harvested from these cells. Purified parasites treated at 12°C for 1 h with freshly prepared 1 mg/mL pronase (Roche)/0.01% saponin/PBS before washing, placing them on slides pre-coated with Cell-Tak (Corning) and fixing with 4% formaldehyde in PBS (20 min, RT). Fixed parasites were washed three times with PBS and permeabilized by incubation with PBS with 0.1% Triton X-100 for 10 min at RT. Slides were then mounted with Mowiol and imaged as above. Samples were coded and enumerated blindly.

### Tachyzoite growth assays

To determine the contribution CPB, CPL and ASP1 on intracellular tachyzoite growth, the relative growth rate of Pru and each knockout strain was compared based on the number of parasites per parasitophorous vacuole 24 and 48 h post-invasion. Freshly harvested tachyzoites were placed onto a monolayer of confluent HFF cells grown on cover slips in a 6-well plate containing DMEM supplemented with 10% CCS. Parasites were allowed to invade for 1 hour, at which point parasites that had not invaded were washed away with DMEM media supplemented with 10% CCS. Parasites were left to grow for a further 24 and 48 h before washing with PBS three times and fixing with 4% (v/v) methanol-free formaldehyde in PBS. Cells were permeablized, blocked and washed as described above. To visualize individual parasites for counting, the prepared slides were incubated for 40 min at RT with rat anti-SAG1 antibodies, which were generated by a commercial service (Pierce) using baculovirus-derived recombinant SAG1 (kindly provided by Martin Boulanger), followed by secondary Alexa-594-conjugated anti-rat antibody (Jackson Immunoresearch). The number of parasites per parasitophorous vacuole was counted for each strain.

### Puncta assays

To assess the formation of puncta (accumulation of dense material) in each strain, we followed the puncta quantification assay described previously (26). Briefly HFF cell monolayers on cover slips were infected with *T. gondii* tachyzoites and differentiated to bradyzoites over the course of 7 days (as described above). For direct immunofluorescence of bradyzoite cyst walls, fluorescein labeled *Dolichos biflorus* agglutinin (DBA; Vector labs) was diluted 1:400 and incubated with infected monolayers that had been fixed and made permeable (as described above). The fluorescent signal derived from DBA staining allows automatic detection of the bradyzoite-containing cyst area using ImageJ software. Cyst images were captured at 100X oil objective as described above. Automatic bradyzoite cyst identification and puncta quantification was performed in ImageJ, using the following parameters described previously (26). Max Entropy thresholding on the GFP channel was used to identify cysts. This was followed by the identification of objects with areas between 130 and 1,900 μm^2^. Particles (puncta) under the GFP mask (therefore within a cyst) were analyzed by automatic local thresholding on the phase image using the Phansalkar method, with the following parameters: radius = 5.00 μm; k value = 0.5; r value = 0.5. Puncta were measured from the resulting binary mask by particle analysis according to the following: size = 0.3–5.00 μm; circularity = 0.50–1.00.

### Bradyzoite Differentiation Assay

Immunofluorescence assays were performed to determine the relative rate at which parasites differentiated from the tachyzoite stage to bradyzoite stage, based on the amount of tachyzoite-specific SAG1 and bradyzoite-specific GFP present within parasitophorous vacuoles/developing cysts. Freshly lysed tachyzoites were placed onto confluent HFF monolayers on coverslips (in 6-well plates) and left to incubate at 37°C, 5% CO_2_ for 24 hours. Following this, the media was removed from each well and alkaline media was then added along with placement in incubators lacking CO_2_ to induce bradyzoite conversion (still maintained at 37°C). Coverslips were fixed and stained with the tachyzoite-specific marker rat anti-SAG1 (1:1000) following 1, 2, 3 and 4 days of differentiation. Strains used in this study express GFP under the bradyzoite-specific LDH2 promoter and this is used as a marker for bradyzoites. Cysts that displayed 50% or more of staining by either rat anti-SAG1 antibodies or GFP were considered positive.

### Bradyzoite viability assays

Bradyzoite viability was assessed by combining plaque assay and quantitative polymerase chain reaction (qPCR) analysis of genome number, as previously described (26). Briefly, HFF cell monolayers were infected with *T. gondii* tachyzoites in 6-well plates. Following host cell invasion tachyzoites underwent differentiation to bradyzoites as described above, resulting in the generation of *in vitro* tissue cysts. Differentiation was carried out over the course of 7 days, replacing the alkaline media daily. Following differentiation, the culture media in each well was replaced with 2 mL Hanks Balanced Salt Solution and cysts were liberated from the infected HFF monolayers by mechanical extrusion, by lifting cells with a cell scraper and syringing several times through 25-gauge needles. Then, 2 mL of pre-warmed 2× pepsin solution (0.026% pepsin in 170 mM NaCl and 60 mM HCl, final concentration) was added to each sample and samples left to incubate at 37°C for 30 min. Reactions were stopped by adding 94 mM Na_2_CO_3_, removing the supernatant after centrifugation at 1,500*g* for 10 min at RT and resuspending pepsin-treated parasites in 1 mL of DMEM without serum. Parasites were enumerated and 1,000 parasites per well were added to 6-well plates containing confluent monolayers of HFFs in D10 media, in triplicate. To allow for the formation of bradyzoite plaques, parasites were left to grow undisturbed for 10 days. After this period, the number of plaques in each well was determined by counting plaques with the use of a light microscope. Five hundred μL of the initial 1 mL of pepsin-treated parasites was used for genomic DNA purification, performed using the DNeasy Blood & Tissue Kit (Qiagen). Genomic DNA was eluted in a final volume of 200 µL. To determine the number of parasite genomes per microliter, 10 µL of each gDNA sample was analyzed by qPCR, in duplicate, using the tubulin primers TUB2.RT.F and TUB2.RT.R (41). Quantitative PCR was performed using Brilliant II SYBR Green QPCR Master Mix (Agilent) and a Stratagene Mx3000PQ-PCR machine. The number of plaques that formed per genome was then calculated. For each experimental replicate the plaques/genome was normalized to the parental control, and this normalized value used for statistical analysis.

## ACKNOWLEDGEMENTS

We thank Aric Schultz for help with the automated puncta analysis, Martin Boulanger for providing recombinant SAG1 and My-Hang (Mae) Huynh for proof-reading the manuscript prior to submission. This work was supported by National Institutes of Health grant R01AI120627 (to VBC and MDC) and a grant from the Stanley Medical Research Institute (to VBC) and by funds from the University of Perugia Fondo Ricerca di Base 2018 program of the Dept. of Chemistry, Biology and Biotechnology (to MDC).

## AUTHOR CONTRIBUTIONS

VBC, MDC and ZD conceived the study. CM, DS and ZD performed most of the experiments and analyzed the data. MDC made and confirmed by PCR PΔcpb, PΔ*asp1* and PΔ*asp1*Δ*cpl* strains. GK performed the ingestion experiments with some assistance from CM and made bradyzoite lysates for immunoblots. CM, DS, ZD, MCD and VBC made the figures. CM, DS, ZD and VBC wrote the paper and all authors revised the paper.

