## Supporting information for "*Toxoplasma* Cathepsin Protease B and Aspartyl Protease 1 play recessive roles in endolysosomal protein digestion during infection"

**A**

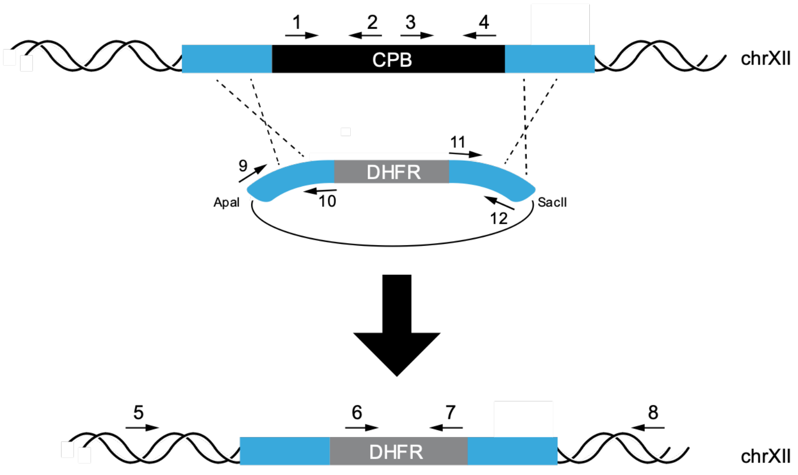


**B**

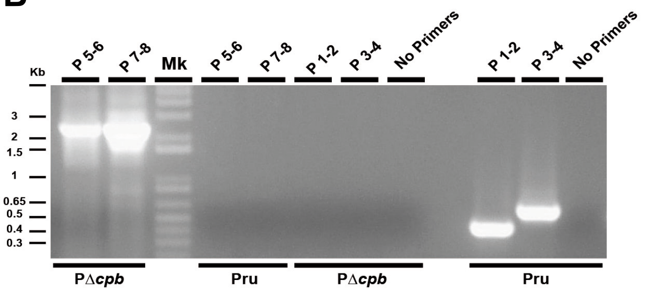


**C**

| Name | Sequence (5' -> 3') |
|---|---|
| P1 | CAGAGTCCGTGGAGTCGTTGTTTGCCAG |
| P2 | GATGAGTGCGACGGCCGCATTCGAGCTG |
| P3 | CTGAAAACGGCGAGGAGTACTGGCATG |
| P4 | CAGGGACTTTACACTCGTTACACAGGAG |
| P5 | GTATTGGGCATGTTCACGTTTCTGTCG |
| P6 | CGCGACGACCAGACACACCGGTTTATGC |
| P7 | CTATCAGTTGTTTTAGTCGAACCGGTTAAC |
| P8 | GGCGCCTAGCACGTTGTGGTTCAGCTC |
| P9 | TACCGGGCCCTTGAGAGGCTACCTAACATCCAG |
| P10 | GGACAAGCTTCTGATGGTGCAGGACTGATATTCCGTC |
| P11 | AGGGGGATCCGTTTGCGGAAAGAGGCGGCTCGAGCAC |
| P12 | TCCACCGCGGGCGGTTCAGCGTATATGAGTGAAC |

**Fig. S1. Creation and validation of PΔ*cpb* parasites.**

**A.** Schematic diagram of double crossover homologous replacement of *CPB* with a dihydrofolate dehydrogenase (DHFR) selectable marker. The vector used to delete *CPB* was generated as follows: 1-kb region of the 5'-UTR of the *CPB* gene was PCR-amplified with *ApaI* and *HindIII* engineered at its 5'- and 3'-ends, respectively, and cloned using these sites in the pMDC64 plasmid upstream of the DHFR cassette carried by this vector. Next, a 1-kb region of the 3'-UTR of the *CPB* gene were amplified by PCR with *BamHI* and *SacII* sites incorporated at its 5'- and 3'-ends, respectively, and cloned downstream to the *DHFR* resistance cassette. The resulting final plasmid was digested with the enzymes *ApaI* and *SacII* to release the backbone from the repair template and introduced into PruΔ*ku80* parasites by electroporation. The transfected parasites were subjected to 2 μM pyrimethamine selection. Clones of the *CPB*-deficient parasites were isolated by limiting dilution. **B.** Replacement of the *CPB* gene with DHFR was confirmed by PCR. **C.** Primers used to generate the *CPB* knockout vector, to test either the integration of DHFR cassette in the *CPB* locus or the absence of the *CPB* gene.

A

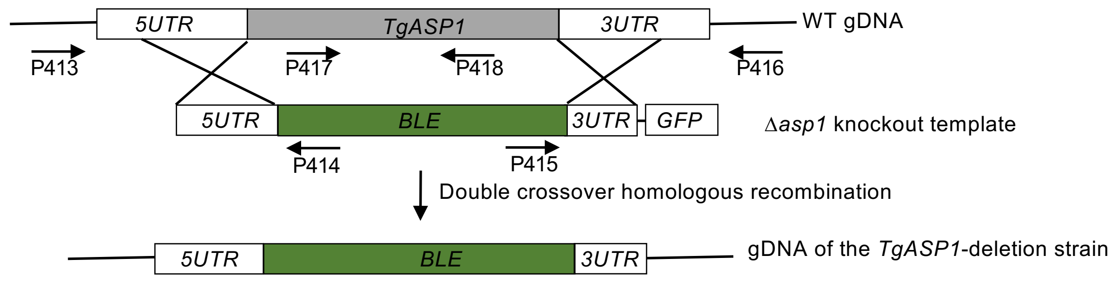


B

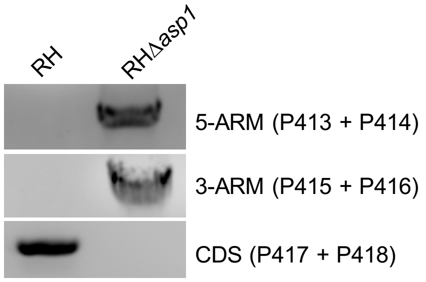


C

| **Primers used to generate the Asp1 KO vector** | **Sequence (5' -> 3')** |
|---|---|
| ASP15UTR.Kpnl.Fwd | CCGGGGTACCGAGAAAAATGGCATGATTGG |
| ASP15UTR.Apal.Rev | AATTGGGCCCCGTGCCCGCGTCCGTAGTTG |
| ASP13UTR.Spel.Fwd | ATCCACTAGTCACAAAGTGCTGCGGTAGTC |
| ASP13UTR.Xball.Rev | ATGCTCTAGAACTAGACGCAGGTCCGGGATAATAAC |
| GFP.Xbal.F | TAGTTCTAGAGCGGCCGCCAC |
| GFP.Xbal.R | ATGC TCTAGA AGTCTTAATTAATTATTTGTATAGTTCATCC |
| **Primers to test integration of selection cassette and Asp1 deletion** | **Sequence (5' -> 3')** |
| P413 | AACGTGGAGACGCCCAGCACC |
| P414 | CCACTCCGCGCGCTTCAGTGG |
| P415 | ACATGGGTCAATACACAAGAC |
| P416 | ACAGACGCGGGAAGGAGAGC |
| P417 | AACTTGCGTCAGTAATCGCTG |
| P418 | TCTACAAAATAGAAGCCATCC |
| **Primers used to generate the Asp1 complement vector** | **Sequence (5' -> 3')** |
| 5UTR.ASP1.F | GCGAATTGGGTACCGGGCCCGGCCTGACTATATGCAACGG |
| 5UTR.ASP1.R | AACCGCGACGACGGAGACATCGTGCCCGCGTCCGTAGTTG |
| ASP1.cds.F | CAACTACGGACGCGGGCACGATGTCTCCGTCGTCGCGGTT |
| ASP1.cds.R | GACTACCGCAGCACTTTGTGTCAGTTCTTGAGTCTGGCGA |
| 3UTR.ASP1.F | TCGCCAGACTCAAGAACTGACACAAAGTGCTGCGGTAGTC |
| 3UTR.ASP1.R | ATACCGTCGACCTCGAGGGGAGTTTAGACGGCCGCTGCAC |

**Fig. S2. Creation and validation of Δ*asp1* and Δ*asp1ASP1* strains.** To generate the TgASP1-deletion mutant, a 3-kb region of the 5’-UTR of the *TgASP1* gene was PCR-amplified with *KpnI* and *ApaI* sites engineered at its 5’- and 3’-ends, respectively. The resulting PCR product was digested by *KpnI* and *ApaI* and cloned in a plasmid upstream to the bleomycin resistance cassette (BLE). Next, a 3-kb region of the 3’-UTR of the *TgASP1* gene was amplified by PCR with *SpeI* and *XbaI* sites incorporated at its 5’- and 3’-ends, respectively, and cloned downstream to the *BLE* resistance cassette. To distinguish random integration of the *BLE* cassette into the parasite genome, a GFP expression cassette was PCR-amplified with *XbaI* sites engineered at both ends and cloned downstream from the 3’-UTR region of *ASP1*. **A.** The resulting final plasmid was introduced into wildtype RH parasites by electroporation. The transfected parasites were subjected to bleomycin selection at 50 μg/ml twice during their extracellular stage. Clones of the *ASP1*-deficient parasites were isolated by limiting dilution. To complement Δ*asp1* mutant, we PCR-amplified the coding sequence of *TgASP1* from the *Toxoplasma* cDNA library and ~1kb of 5’- and 3’-UTRs of *TgASP1* from *Toxoplasma* genomic DNA. All three PCR fragments were gel-purified and assembled into the pMDC64 plasmid which encodes a pyrimethamine resistance cassette by Gibson Assembly. The resulting ASP1 complementation construct was electroporated into the Δ*asp1* parasites, followed by pyrimethamine selection and limited dilution to isolate single clones. **B.** Removal of the *ASP1* gene by BLE cassette was confirmed by PCR. **C.** Primers used to generate the *ASp1* knockout vector, to test either the integration of BLE cassette in the *ASP1* locus or the absence of the *ASP1* gene.

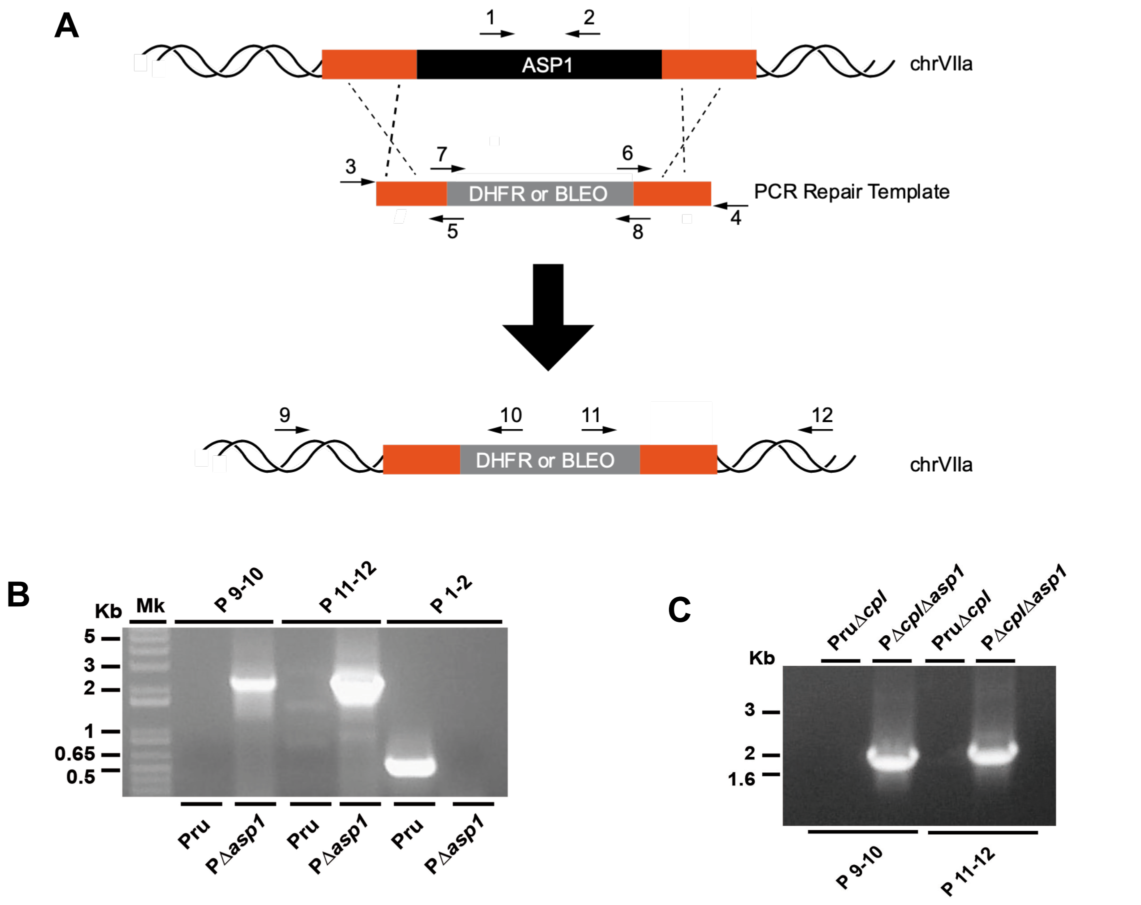


D

| Name | Sequ ence (5' -> 3') |
|---|---|
| P1 | CTGCAACTTGAGATCGAAGGGCAGGACTA |
| P2 | AAGGCTTTCAGGCCCATCATGCCGAAGA |
| P3 | GAAAGCTCTAGGGGCCACCAGTGTCACT |
| P4 | ACCCATGAATGCAATCGCTAACGCCTAC |
| P5 | CGCTGATCAGAGAATTTCACGAACGCGGACAAGCTTATCGATACCGTCGA CCTCGGTAGTTGTAGGACTAATAGACACTCGA |
| P6 | AACACAATTCTTGCCCCCCCGAGGGGGATCCACTAGTTCTAGAGCGGCCGC CACCGCACACCGAATAAGACGGCACTTACAC |
| P7 | TCGAGTGTCTATTAGTCCTACAACTACCGAGGTCGACGGTATCGATAAGCT TGTCCGCGTTCGTGAAATTCTCTGATCAGCG |
| P8 | GTAAGTGCCGTCTTATTCGGTGTGCGGTGGCGGCCGCTCTAGAACTAGTGG ATCCCCCTCGGGGGGGCAAGAATTGTGTTAACC |
| P9 | GTAGGTTTTAGGTACACGCAGACCTCGCA |
| P10 | TCTGCGATGAGCGCAAGCCGTCCTGA |
| P11 | CACCGTTGTGCTCACTTCTCAAATCGAC |
| P12 | CTTCCAGATAATCTCAGTTGTCTGTGAGT |

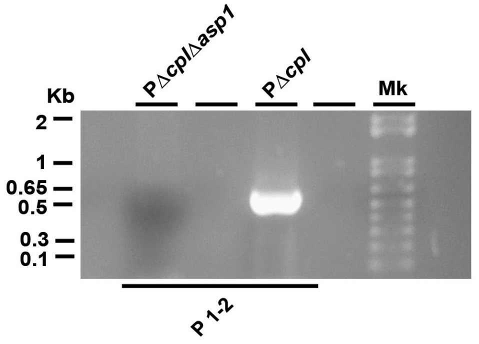


**Fig. S3. Creation and validation of PΔ*asp1 and* PΔ*cpl*Δ*asp1* strains. A.** Schematic diagrams of homologous replacement of *ASP11* with either DHFR or BLE selectable marker in PruΔ*ku80* and PΔ*cpl*, respectively. The repair template used to obtain *ASP1* deletion was generated as follows: in a first run of PCR, three DNA fragments encompassing 1-kb region of the ASP1 5'-UTR, the selectable marker and the *ASP1* 3'-UTR were generated. The selectable marker was amplified using a forward and reverse primer carrying 40 bp complementary to the ASP1 5'- and 3'-UTR, respectively, to allow the fusion of the 3 fragments during a second round of PCR using the forward and reverse primer binding to the ASP1 5'- and 3'-UTR, respectively. Five μg of the fusion PCR product was introduced into either PruΔ*ku80* or PΔ*cpl* parasites by electroporation. The transfected parasites were subjected to drug selection and *ASP1*-deficient clones were isolated by limiting dilution. Deletion of the *ASP1* gene was confirmed by PCR in PruΔ*asp1* (**B**) and *P*Δ*cpl*Δ*asp1* (**C**). **D.** Primers used to generate the *asp1* KO repair template and to test either the integration of selectable cassette in the *asp1* locus or the absence of the asp1 gene.
